# Non-invasive Stimulation of Contralateral Primary Motor Cortex Reduces the Amount of Skill Generalization to the Untrained Arm

**DOI:** 10.1101/2025.07.22.666199

**Authors:** Goldy Yadav, Manon Chauvaux, Julie Duque

## Abstract

Successfully learned motor skills can generalize or transfer to the untrained arm. The neural substrate underlying such intermanual/interlimb generalization of newly acquired skill memory is unclear. Here, we focused on contralateral primary motor cortex (cM1) which is considered a key brain area for skill learning and memory consolidation. We probed the causal role of cM1 in intermanual skill generalization in a two-day study involving right-handed young individuals (n=31) who learned a novel motor skill reaching task. Immediately following (right-arm) learning, we delivered low-frequency (1Hz, 1800 pulses) repetitive transcranial magnetic stimulation (rTMS) to target left cM1 in one group of individuals (n=15), while another group (n=16) served as an active control in which ipsilateral M1 (iM1) was targeted. On the same day we measured corticospinal excitability (CSE) to assess learning-induced and rTMS-induced neuroplastic changes occurring in the targeted M1s. Next day after 24-hours, both groups were tested for intermanual skill generalization (left-arm), followed by a brief test of intralimb retention (right-arm). Our results show that stimulating cM1, versus iM1, reduced the amount of generalization to the untrained arm on the next day, without affecting its (re)learning ability or the follow-up retention performance of the trained arm. Further, rTMS stimulation induced a net facilitation in CSE- with higher facilitation tending to correlate to lower generalization in a subset of high learners in cM1 group. Taken together, this study highlights the role of cM1 in skill generalization such that it seems to mediate the early transfer of learning to the untrained arm.

**NEW AND NOTEWORTHY:** Intermanual skill generalization from trained to the untrained arm is causally mediated by the contralateral (trained) primary motor cortex (cM1) as opposed to the ipsilateral (untrained) motor cortex. Low-frequency stimulation of M1 in our study led to a facilitation of corticospinal excitability, while impairing the amount of skill generalization in the cM1 group. This highlights a rather paradoxical and opposite effect of non-invasive brain stimulation, such as rTMS, on motor behavior and associated motor excitability. Our data also suggest involvement of additional brain areas for such motor skill behavior that were rendered unperturbed by rTMS in this case and, thus, contributed to an overall positive skill performance achieved by both arms the next day.

## 1. INTRODUCTION

Human motor skill behavior is dependent on formation and stabilization of motor memories resulting from successful learning. Such motor memories are deemed robust and especially beneficial when they can generalize to untrained contexts, for instance the untrained arm. This form of generalization is known as intermanual generalization (or interlimb transfer) and, given its relevance for clinical rehabilitation, it is a widely studied topic in the domain of motor neuroscience (Ghahramani et al., 1996; Krakauer et al., 2000; Hwang et al., 2003; Shadmehr, 2004; Farthing et al., 2009; Magnus et al., 2010; Ausenda and Carnovali, 2011; Pearce et al., 2013; Green and Gabriel, 2018; Kumar et al., 2020; Yadav and Mutha, 2020; Yadav et al., 2025a). In the domain of reaching movements, most research studies have shown that a newly learned reaching movement can generalize to the untrained arm in healthy individuals. Yet, these studies have used different motor learning paradigms leading to quite heterogenous findings when it comes to characterizing such interlimb generalization. For example, studies using motor adaptation paradigms have shown that such generalization is mostly unidirectional or asymmetrical i.e., left arm to the right arm only (Wang and Sainburg, 2003, 2004; Bao et al., 2017; Kumar et al., 2018; Kumar et al., 2020), and can be influenced by factors such as training speed (Lefumat et al., 2015) and contextual similarity of the training and task space (Hwang et al., 2003; Wang and Sainburg, 2006). Other studies using non-adaptation paradigms such as training for ballistic movements (Hinder et al., 2011; Hinder et al., 2013; Stöckel et al., 2016) have reported that interlimb generalization can occur from right to left arm and vice versa. Finally, our work focusing on de novo skill learning showed that newly acquired motor skills characterized by improved movement speed and accuracy (Yadav and Duque, 2023) can generalize from dominant right arm to the left non-dominant arm following long training (Yadav et al., 2025a); as well as from non-dominant to the dominant arm (Yadav and Mutha, 2020).

In contrast to this significant amount of research work exploring behavioural aspects of interlimb generalization of reaching movements, the neural basis of this generalization of newly learned motor skills is unclear. The current paper aims to address this gap in knowledge and, as a first step, we focus on elucidating the role of primary motor cortex (M1) as the neural substrate mediating interlimb skill generalization. It is widely known that M1 is a critical brain region in the network that drives motor skill learning (Floyer-Lea and Matthews, 2005; Halsband and Lange, 2006; Doyon et al., 2009; Dayan and Cohen, 2011; Cantarero et al., 2013; Narayana et al., 2014; Kawai et al., 2015). More specifically, studies have shown that damage to M1 can lead to pronounced contralesional impairments of fine motor skills (Canning et al., 2000; Balasubramanian, 2015; Buetefisch et al., 2018; Gerardin et al., 2024). Further, non-invasive stimulation of the contralateral M1 (cM1) in healthy individuals has been shown to enhance online and offline skill performances (Reis et al., 2009) as well as learning related changes in motor excitability (Yadav et al. 2025b). On the other hand, repetitive transcranial magnetic stimulation (rTMS) over cM1 can disrupt skill acquisition (Kobayashi et al., 2009). In addition, Kantak et al., (2010a) showed the causal involvement of cM1 in memory consolidation of a newly learned skill (skilled reaching movements to predictable target) such that rTMS over cM1 immediately after training impaired retest performance of the trained arm at 24 hours. Parallelly, there is converging evidence for cM1 involvement in enhancing memory of newly learned motor sequences (Hussain et al., 2021) as well as value-based motor decisions (Derosiere et al., 2017a; 2017b).

Taken together, it is evident that cM1 undergoes significant plastic changes during skill training and supports memory formation and consolidation processes that can influence future motor performances. On the other hand, as mentioned earlier, intermanual skill generalization is contingent on factors that influence training arm performance and processes that develop during training. Thus, it is possible that training induced neuroplastic changes occurring within cM1 also drive generalization of the learned skill to the untrained arm. To causally probe this idea, we recruited 34 young healthy individuals (31 included in the final analysis) who trained on a novel motor skill in a predictable training condition (known to specifically engage cM1) with their right (dominant) arm. Immediately following training, we applied rTMS to disrupt M1, contralateral to the trained arm in the test group (cM1 Group-rTMS over left M1) or ipsilateral to the trained arm in an active control group (iM1 Group-rTMS over right M1). Finally, we tested the untrained left arm of all individuals 24-hour later for intermanual skill generalization. Then, we also tested the trained right arm for intramanual skill retention performance to assess if rTMS to M1 following training on Day-1 also impaired follow-up retention performance on Day-2. We expected that rTMS to cM1, as compared to iM1, would result in significant impairments in intermanual skill generalization, and possibly also in intramanual retention at 24 hours because such non-invasive stimulation of cM1 can disrupt memory representations starting to consolidate immediately following skill training.

## 2. MATERIALS AND METHODS

Thirty-one (out of the recruited 34, see Statistical Analyses section below) right-handed young individuals (25 Females; 23.09 ± 2.80 years old; normal or corrected vision) successfully completed the entire study. The current study spanned over two consecutive days and involved behavioral assessments (motor skill learning, generalization and retention) and neurophysiological procedures involving transcranial magnetic stimulation (TMS) and electromyography (EMG) recording. Prior to participating in the study, all individuals completed a handedness questionnaire (adapted from Hull, 1936), as well as a TMS risk assessment questionnaire to rule out any contraindications. None of the participants reported any neurological or psychiatric disorder or had any history of drug/alcohol abuse. All Individuals provided written informed consent and were financially compensated for participating in the study. The study was conducted in accordance with the Ethics Committee of the University and the principles of the Declaration of Helsinki.

### 2.1 Experimental Design

#### 2.1.1 Motor skill task

##### Apparatus

The task setup comprised of a graphic tablet (Huion, Graphics Technology (HK) Limited) placed horizontally on a tabletop connected to a vertical computer screen (Figure 1A). Participants were required to make planar reaching movements on the tablet with a stylus affixed to their index finger which provided real-time position coordinates data during the movement. Finger position data were sampled at 60Hz. The task was displayed on the computer screen and designed using MATLAB with Psychtoolbox (Brainard and Vision, 1997).

**Figure 1.**
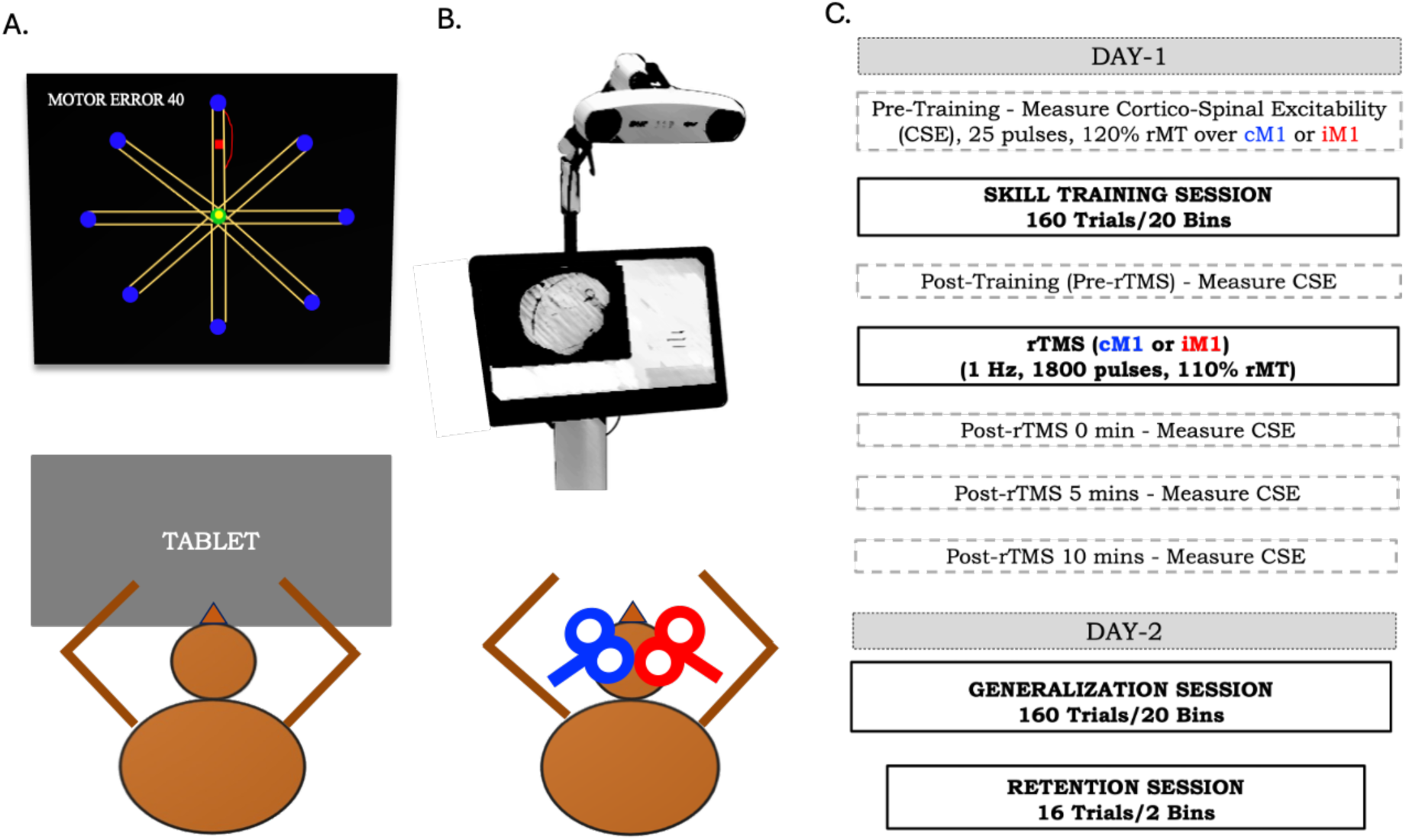
Experimental Design. **A) Motor skill task** in which right-handed participants (n=31) were required to make fast and accurate movements on a digitizing tablet to one of the eight target circles (blue color) presented clockwise on a computer screen (shown on top) with a goal to reduce ‘Motor Error’ (numeric score computed based on movement time and accuracy). **B) TMS procedures** over left/contralateral M1 (test group cM1 in blue, n=15) or the right/ipsilateral M1 (active control group iM1 in red, n=16)) were performed using Neuronavigation (shown on top). **C) Study design flow** depicted for Day-1 (with TMS procedures and skill training session) and Day-2 (skill generalization and retention test sessions). Note that right arm was used for skill training as well as for retention test, while left arm was tested during generalization session in all participants.

##### Task Procedure

The skill task (a modified version of skill task used in Yadav et al., 2025a) started with positioning the hand inside a green start circle (1cm diameter) at the center of the tablet. After 500 ms of staying within the start circle, a blue target circle (1 cm diameter) appeared on the screen (at a distance of 15 cm) along with an audio ‘go’ cue. Subjects were instructed to make fast and accurate movements from the start position (green circle on the screen) to the target (blue) circle in 750 ms, within a specified path- a 1 cm wide yellow path connecting the green and blue circles (Figure 1A). Performance feedback was given at the end of each trial-temporal error was indicated by a red square denoting hand position at 750 ms after movement onset, while spatial error was indicated by a red line displaying parts of the movement falling outside the specified yellow path. In addition to these visual cues (see Figure 1A), participants were also provided with a numeric score indicating *‘Motor Error’* (in mm) which was a sum of temporal and spatial errors (a measure used in our previous work of Yadav and Mutha, 2016; 2020 and Yadav et al., 2025a; and detailed later in section 2.2.1). Participants were instructed to reduce this motor error over the course of training by performing fast and accurate movements to the target circle. On each trial, one target circle appeared at one of the eight locations on the screen (target location was clockwise in order, and thus predictable). Each trial lasted 2 seconds (movement time), and the performance feedback (visual cues with numeric motor error score) were presented for an additional 2 seconds on the screen.

All participants first received a brief familiarization on the motor skill task (8 trials each with right and left arm, respectively). Next, during the ‘*Skill Training Session’* on Day-1, they performed a total of 160 trials (8 trials corresponding to the eight target directions and repeated over 20 blocks) to learn the skill task with the right hand. Immediately, after training on that day, all subjects received repetitive TMS (rTMS) over M1 (see Figure 1B and following section for more details). Next, on Day-2 during the ‘*Generalization Session’*, all subjects performed 160 trials (20 blocks) with their left untrained hand to test intermanual skill generalization. Immediately following this on Day-2, we also briefly had subjects perform 16 trials (2 blocks) with their trained right hand to evaluate intramanual skill retention performance at 24 hours (*‘Retention Session’*, see Figure 1C).

#### 2.1.2 Transcranial Magnetic Stimulation (TMS)

##### TMS Procedure with Neuronavigation

As mentioned in the introduction, our main objective was to assess the causal role of contralateral (left) M1 in interlimb generalization of newly learned motor skill in young right-handed individuals. To probe this brain region, we used a figure-of-eight TMS coil (MCF-B70 connected to MagPro X100 stimulator, Magventure) along with a neuronavigation system (visor2™ ANT Neuro). Participants were randomly divided into two groups in the beginning of the experiment on Day-1. In one group, TMS pulses were delivered to the region of interest i.e., left contralateral M1 (cM1 Group, n=15) while in the second group, TMS pulses were delivered to the right ipsilateral M1 (iM1 Group, n=16) which served as an active control in this study (Figure 1B).

Participants wore a fitted headcap (Electro-cap, Electro-Cap International, USA) onto which neuronavigation trackers were attached to precisely co-register head position with respect to the TMS coil. TMS coil was placed tangentially at 45-degree angle from the midline on the scalp. TMS pulses were delivered over cM1 or iM1 (depending on the group) to identify the optimal scalp position where a single pulse of TMS consistently produced the best detectable motor-evoked potential (MEP, recoded using EMG detailed in next section) in the contralateral first dorsal interosseous (FDI of the right or left hand, respectively), an agonist muscle for our skill task. This “hot spot” was registered in the neuronavigation system for each participant (onto a default MRI) and used as reference throughout the experiment. Next, we proceeded to determine resting Motor Threshold (rMT) at the hotspot for each participant, which refers to the minimal TMS intensity required to evoke MEPs of ∼50 µV peak-to-peak in the target muscle (right or left FDI) in at least 5 out of 10 consecutive pulses.

TMS was used on Day-1 only. The hotspot and rMT were established before the *Skill Training Session* in all individuals of both groups (mean rMT of group cM1=53.33%; iM1= 53.17% of maximum stimulator output). Our study design also allowed us to probe changes in corticospinal excitability (CSE) as a consequence of training on our skill task. To do so, we applied single biphasic TMS pulses on the registered M1 hotspot (cM1/iM1 depending on the group) at an intensity of 120% of the individual’s rMT. Twenty-five pulses with inter-pulse interval of 4 - 4.3 seconds were delivered to obtain reliable MEPs in the right (trained hand for cM1 group) or left (untrained hand for iM1) FDI muscle (depending on the group) before and after the *Skill Training Session* (see Figure 1C). Next, central to our research question, we applied rTMS to probe the causal role of cM1 in interlimb generalization of motor skill learning. To do so, we employed a widely studied low frequency rTMS (1Hz) neuromodulation protocol, considered to have an inhibitory effect on the stimulated region possibly via long-term depression (LTD) like mechanisms (Hess and Donoghue, 1996; Chen et al., 1997; Boroojerdi et al., 2001; Fitzgerald et al., 2002; Kantak et al., 2010a, 2010b). We delivered 1800 pulses of 1Hz rTMS (lasting about 30mins) at an intensity of 110% of rMT. This rTMS intervention was applied over cM1 (test group) or iM1 (control group). To assess rTMS-induced changes in CSE associated with this neuromodulation intervention, we delivered single-pulse TMS to obtain resting MEPs immediately before, as well as at three different time points after the intervention- at 0, 5 and 10-mins following rTMS offset. As mentioned earlier, 25 pulses (inter-pulse interval of 4 - 4.3 seconds at 120% of rMT) were delivered again to obtain the resting MEPs at each of these time points following rTMS.

##### Electromyography (EMG) Recording and Processing

EMG was recorded to measure peak-to-peak MEP amplitude using surface bipolar electrodes targeting the FDI-one electrode was placed on the belly of the muscle; a second electrode was placed on the proximal interphalangeal joint of the index finger and a ground electrode was placed on the ulnar styloid of the wrist. Note that EMG electrodes were attached to both left and right hands of all individuals (regardless of the group) to avoid any confounding effect of electrode placement on skill performance which was always with the right arm only. In other words, this ensured that both cM1 and iM1 groups performed the skill task with electrodes placed over their right training hand. However, EMG was recorded and assessed from one hand (FDI) only, depending on the group type (right hand for cM1 and left hand for iM1 group). The raw EMG signal recoded from the target FDI muscle was amplified (1K gain), band-pass filtered (10-500 Hz, Neurolog; Digitimer), notch-filtered (50 Hz) (Digitimer D360, Hertfordshire, UK) and digitized at a sampling rate of 2 kHz (CED and Signal 6 software) for off-line analysis (similar to Yadav et al., 2025b). Data extraction and offline analyses were performed using Signal 6 and MATLAB (Mathworks) software.

### 2.2 Statistical Analyses

For statistical analyses, we used JMP software (Version Pro 17.1, SAS Institute Inc.). For all statistical tests significance level was set at 0.05 and post-hoc Tukey test was conducted following a significant main effect in the ANOVA. Partial eta-squared (η^2^p) measure for effect size reported along the test results with data expressed as mean ± standard error (SE) throughout the manuscript.

#### 2.2.1 Motor skill performance assessments: Learning, Interlimb Generalization and Intralimb Retention

As described above in section 2.1, participants were required to learn a reaching skill task by making fast and accurate movements to reduce the motor error (in mm) during the training session (comprising 20 blocks of 8-trial each). Trials on which participants failed to initiate a movement from the start circle in 750 ms (expected time to reach the target circle, as mentioned earlier) were excluded from our analyses (2.23% of the total trials).

To quantify learning on this task in both groups on Day-1, we assessed change in their mean motor errors over 20 blocks of the training session. Motor error was the main variable of interest for assessing skill performance in this study-sum of temporal error [measured as distance (in mm) of the hand at 750 ms from the target circle] and spatial error [measured as distance (in mm) of the hand path falling outside the specified path connecting the start and target circles]). Participants who exhibited a reduction in the motor error on block-20 as compared to block-1 were included for the remainder of data analyses in this study (a total of 31 participants out of the 34 that were recruited). Next, related to the central question of our study, we statistically examined and compared the extent of skill generalization on Day-2 following skill training and rTMS intervention on Day-1 in both groups. To do so, we performed a two-way repeated-measures ANOVA with group (cM1, iM1) and block (first block training, last block training, first block generalization, last block generalization) as between-subject and within-subject factors, respectively to compare the motor errors. To assess whether intramanual retention performance of the trained right arm on Day-2 was impacted by rTMS following end of training on Day-1, we performed a two-way ANOVA with group (cM1, iM1) and block (last block training, first block retention, last block retention) as factors to compare the motor errors.

#### 2.2.2 CSE measurements in relation to motor skill learning and rTMS intervention

We extracted Root Mean Square (RMS) of EMG activity of the FDI muscle in the 200 ms time window preceding a TMS pulse and peak-to-peak MEP amplitude in the 10-60 ms time window after the TMS pulse using Signal software. This was done to obtain resting MEPs corresponding to the five time points of CSE measurement (see Figure 1C), i.e., Pre-Training, Post-Training (also Pre-rTMS), Post-rTMS 0 min, Post-rTMS 5 min and Post-rTMS 10 min. We removed the first MEP of each time point, as well as any MEP preceded by a significant muscular activity (based on RMS threshold set at 0.02 mV), as done previously (see Methods section of Yadav et al., 2025b). Next, we removed outlying trials (± 2.5 SD of mean of each time point), as also typically done in our lab (see for instance, Quoilin et al., 2019). As a result, a total of 6.4% of all trials were removed, leaving at least 21 MEPs (out of 25 pulses delivered) for each TMS time point for each participant. Finally, we calculated the mean MEP amplitude for each of these five TMS time points.

First, we assessed the consequence of de novo motor skill learning on CSE by analyzing MEPs with a two-way ANOVA with group (cM1, iM1) and TMS timing (Pre-Training, Post-Training) as factors. Remember that cM1 MEPs were recorded from the trained right arm, whereas iM1 MEPs were recorded from the untrained left arm. Therefore, the factor ‘group’ in this analysis reflects differences in CSE not only across subject groups, but also across different arms with respect to training. Then, in a further analysis, we linked training-related CSE changes i.e., ΔCSE_training_ (Post-Training minus Pre-Training MEPs) to motor error changes (First-Training minus Last-Training blocks) to explore any association between the amount of skill learning on our task and underlying CSE changes. A positive correlation would indicate that a greater training-induced CSE increase is associated with greater learning. Finally, we assessed the effect of the 1Hz rTMS on CSE. To do so, we compared MEPs obtained after the training session, but immediately before the rTMS intervention (in other words, pre-rTMS), with post-rTMS (0, 5, 10 mins) time points (see Figure 1C). To compare MEPs obtained at these four different time points, we performed a two-way ANOVA with group (cM1, iM1) and rTMS timing (Pre, Post-0 min, Post-5 min, Post-10 mins) as factors. Based on this finding in which we observed net facilitation in CSE, we performed further correlations to explore any underlying relationship between CSE changes following rTMS i.e., ΔCSE_rTMS_ (Post-rTMS 10 min minus Pre-rTMS MEPs) and intermanual generalization performance i.e., ΔGeneralization (Motor Errors on First-Training minus First-Generalization blocks) of our test group participants (cM1). Here, a negative correlation would indicate that a greater rTMS-induced CSE facilitation is followed by an attenuated generalization on the next day.

## 3. RESULTS

### 3.1 Skill Behavior

#### Learning and Intermanual Generalization

The overall training, generalization and retention performances of cM1 and iM1 groups are depicted in Figure 2. As evident from the left panel of this figure, both groups learned the skill task on Day-1 by reducing their motor errors over the 20 blocks of training session. We, therefore, proceeded to statistically examine the intermanual skill generalization in these two groups resulting from the skill training and rTMS procedure (see Figure 3). Our two-way ANOVA revealed a significant effect of block (F_3,87_ = 117.8091, p<0.0001, η^2^_p_=0.8024) and a block X group interaction (F_3,87_ = 3.0002, p=0.0349, η^2^_p_=0.0937) with no main effect of group (F_1,29_ = 1.1076, p=0.3013, η^2^_p_=0.0755). As expected, Tukey post hoc tests revealed significant differences in mean motor errors on the first and last blocks of the training session on Day-1 in both cM1 (p<0.0001) and iM1 (p<0.0001) groups. We observed that the mean motor errors reduced in both groups over the first (cM1: 93.7 ± 3.69 mm; iM1: 83.7 ± 3.57 mm) and last training blocks (cM1: 30.5 ± 3.69 mm; iM1: 30.4 ± 3.57 mm), with no significant difference between the groups on the first (p=0.5273) or last training (p=1.00) blocks. This suggests that both groups were able to successfully learn the motor skill task in our study with lower motor errors evident by the end of the training session on Day-1 (see Figure 2 and Figure 3 Left Panels).

**Figure 2.**
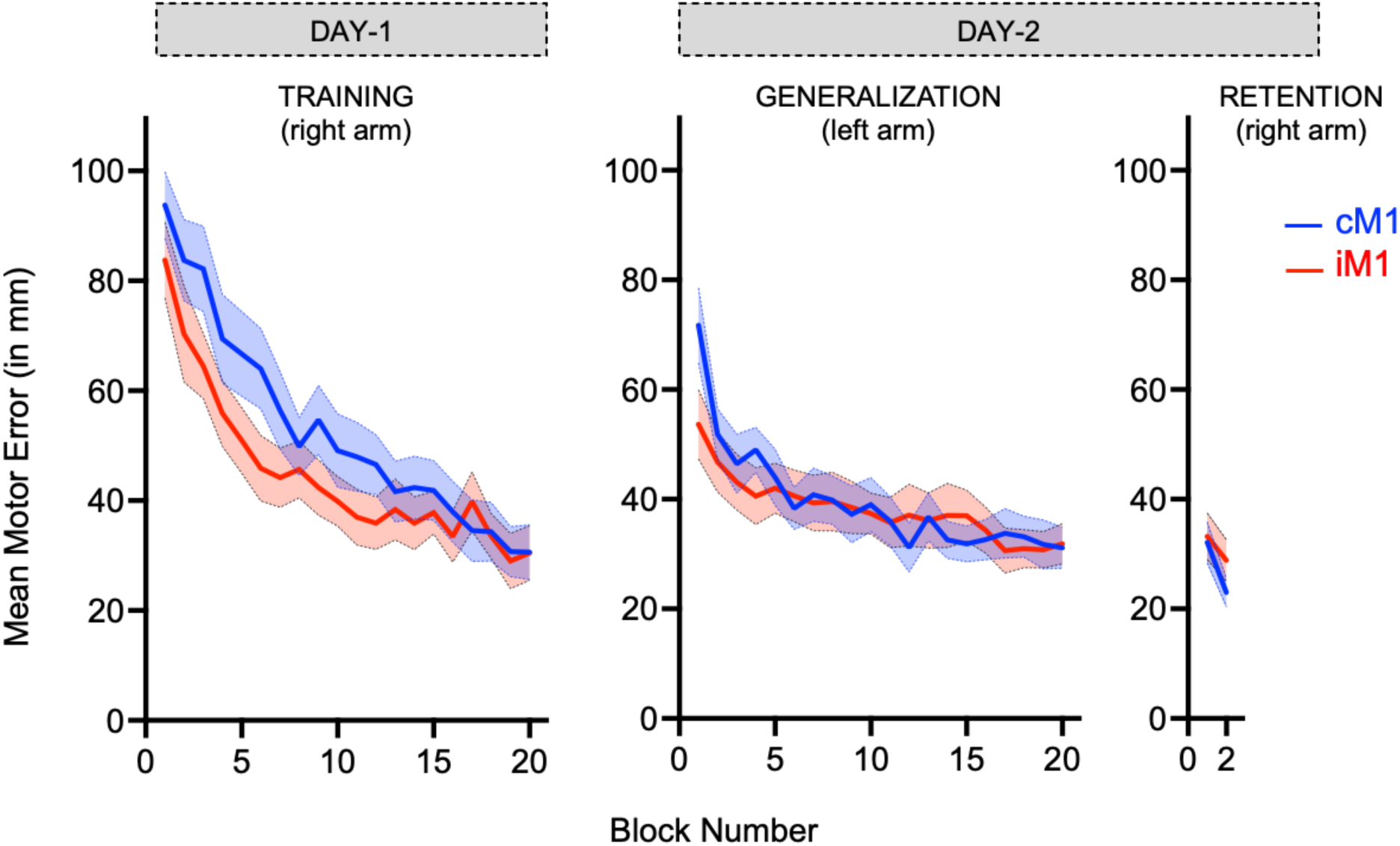
Mean motor error across training, generalization and retention sessions. Left Panel: Skill performance on Day-1. Both groups (cM1 in blue, n=15; iM1 in red, n=16), learned the motor task with their right arm by reducing mean motor errors (in mm) over the 20 blocks during the training session. Middle Panel: Generalization session on Day-2. cM1 group started with larger motor errors as compared to iM1 group, on the first block of generalization session with their untrained left arm. Right Panel: Ensuing retention session on Day-2. Comparable skill performance observed in the two groups when tested for intramanual retention with their trained right arm. Shaded error bands on the figure denote SE.

**Figure 3.**
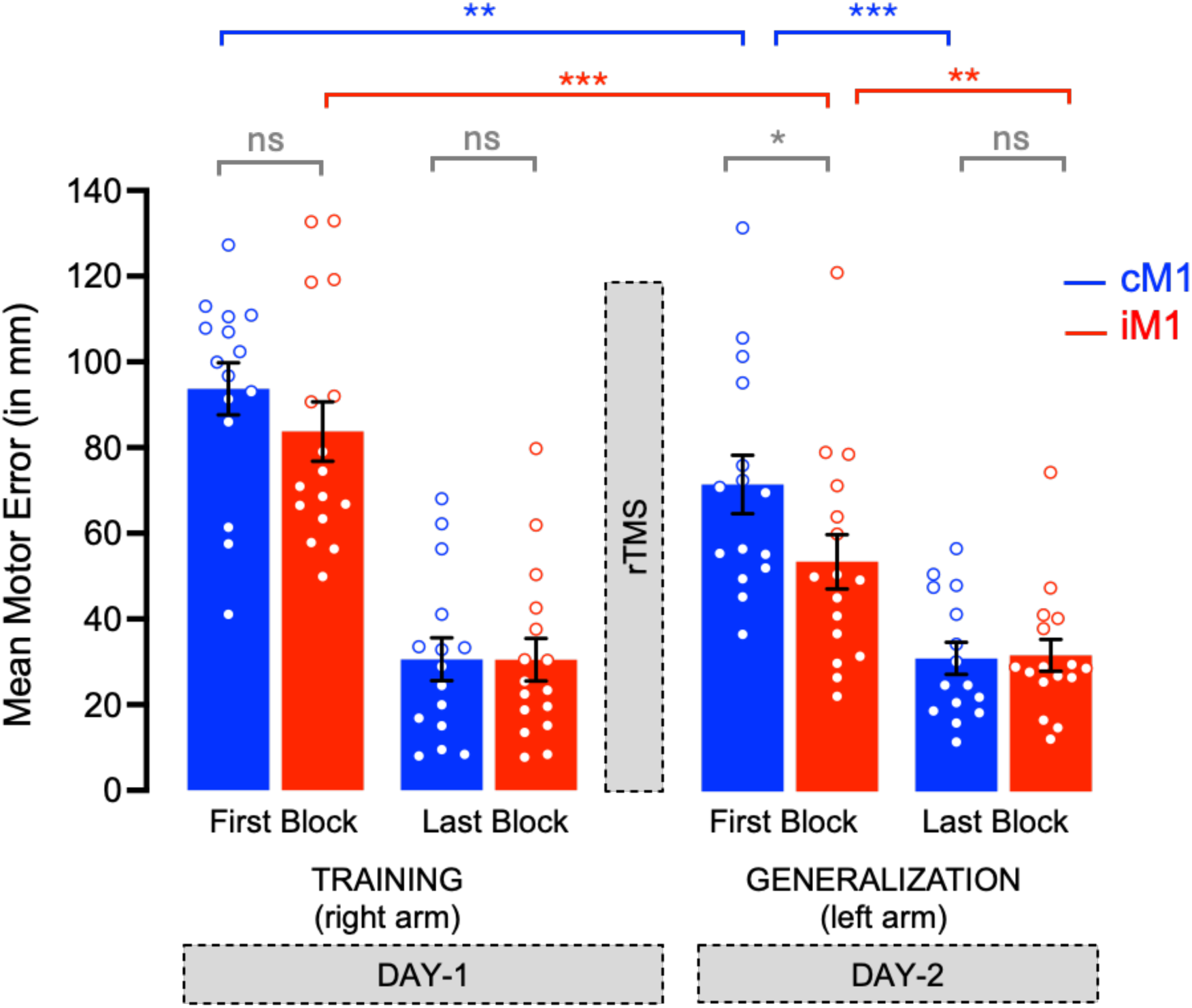
Intermanual Skill Generalization. Left Panel: Lower motor errors observed in both cM1 (in blue, n=15) and iM1 (in red, n=16) groups at the end of Day-1 suggesting comparable level of skill learning. Right Panel: When tested on Day-2, both groups exhibited reduced motor errors on the first generalization block as compared to first training block; a signature of intermanual skill generalization. However, cM1 group (who received rTMS over contralateral M1 at the end of training) exhibited significantly larger motor errors on the first block of generalization session, as compared to iM1 group (who received rTMS over ipsilateral M1 at the end of training), indicating reduced amount of skill generalization in this group. Both groups further improved performance of their left untrained arm and reached a comparable level by the last block of this session. Error bars on the figure denote SE. Colored circles denote individual participants. [*** denotes p<0.0001; ** denotes p<0.001, * denotes p<0.05, ns denotes non-significant]

Next, upon examining intermanual generalization by comparing first training (on Day-1) and first generalization blocks (on Day-2, as we have done previously in Yadav et al., 2020; 2025a), the post hoc test revealed a significant difference in motor errors between these blocks corresponding to the right/trained and the left/untrained arm in both cM1 (p=0.0015) and iM1 (p<0.0001) groups, respectively. Interestingly, however, a direct comparison of motor errors on the first block of generalization (see Figure 2 Middle Panel and Figure 3 Right Panel) between cM1 and iM1 revealed a significant difference between the two groups (p=0.0157) with higher motor errors observed in cM1 group (71.7 ± 3.69 mm) as compared to iM1 (53.6 ± 3.57 mm). These findings indicate that while both groups exhibited significant skill generalization to their untrained arm (on Day-2) thus, benefiting from training of the first arm (on Day-1); it was less prominent in the cM1 group. This suggests that rTMS over cM1 following skill training may have caused this reduced amount of skill generalization observed on the first block of generalization session in this group compared to the iM1 group, as we had hypothesized in the introduction of this paper. However, this impairing effect of rTMS on skill generalization in the cM1 group didn’t last for the entire session as we did not find any significant difference between the two groups on the last block of the generalization (p=1.0000 [cM1: 31.1 ± 3.69 mm and iM1: 31.8 ± 3.57 mm]). We found that the untrained left arm exhibited further significant reduction in the motor errors over the first and last generalization blocks in both cM1 (p<0.0001) and iM1 (p=0.0011) groups. This reflects (re)learning by the untrained arm on Day-2 reaching a performance level that was comparable to the training arm performance at the end of Day-1 (last training and last generalization blocks not significantly different for cM1 (p=1.00) or iM1 (p=1.00).

Overall, these results suggest that a newly learned skill task in our study was generalized to the untrained arm when tested on Day-2 in the two groups. However, the amount of skill generalization (as evident on the first block of generalization test) was reduced in the group that received rTMS over the contralateral M1 immediately following training on Day-1. Further, this rTMS induced impairment was possibly quashed by relearning processes that led to overlapping performances observed in the two groups by the end of the generalization session.

#### Intramanual Retention after 24 hours on Day-2

Finally, related to the intramanual retention session, we assessed and compared the trained right arm performances of the two groups at the end of training (just prior to rTMS intervention on Day-1) with that on Day-2 (which immediately followed the generalization session, see Figure 4). Our group (cM1, iM1) X block (last training, first retention, last retention) ANOVA revealed a significant effect of block (F_2,58_ = 4.5351, p=0.0148, η^2^_p_=0.1352) only, with no group (F_1,29_ = 0.1868, p=0.6688, η^2^_p_=0.0250) or group X block interaction (F_2,58_ = 0.9645, p=0.3872, η^2^_p_=0.0321). Tukey post hoc tests revealed that the only significant difference for motor errors was that between the first and last retention blocks (p=0.0127), regardless of the group. Errors were comparable between the first retention and last training blocks (p=0.6245) as well as between the last retention and last training blocks (p=0.1172). This suggests that rTMS intervention following training on Day-1 did not cause any group differences in retention performance of the trained arms when tested on Day-2.

**Figure 4.**
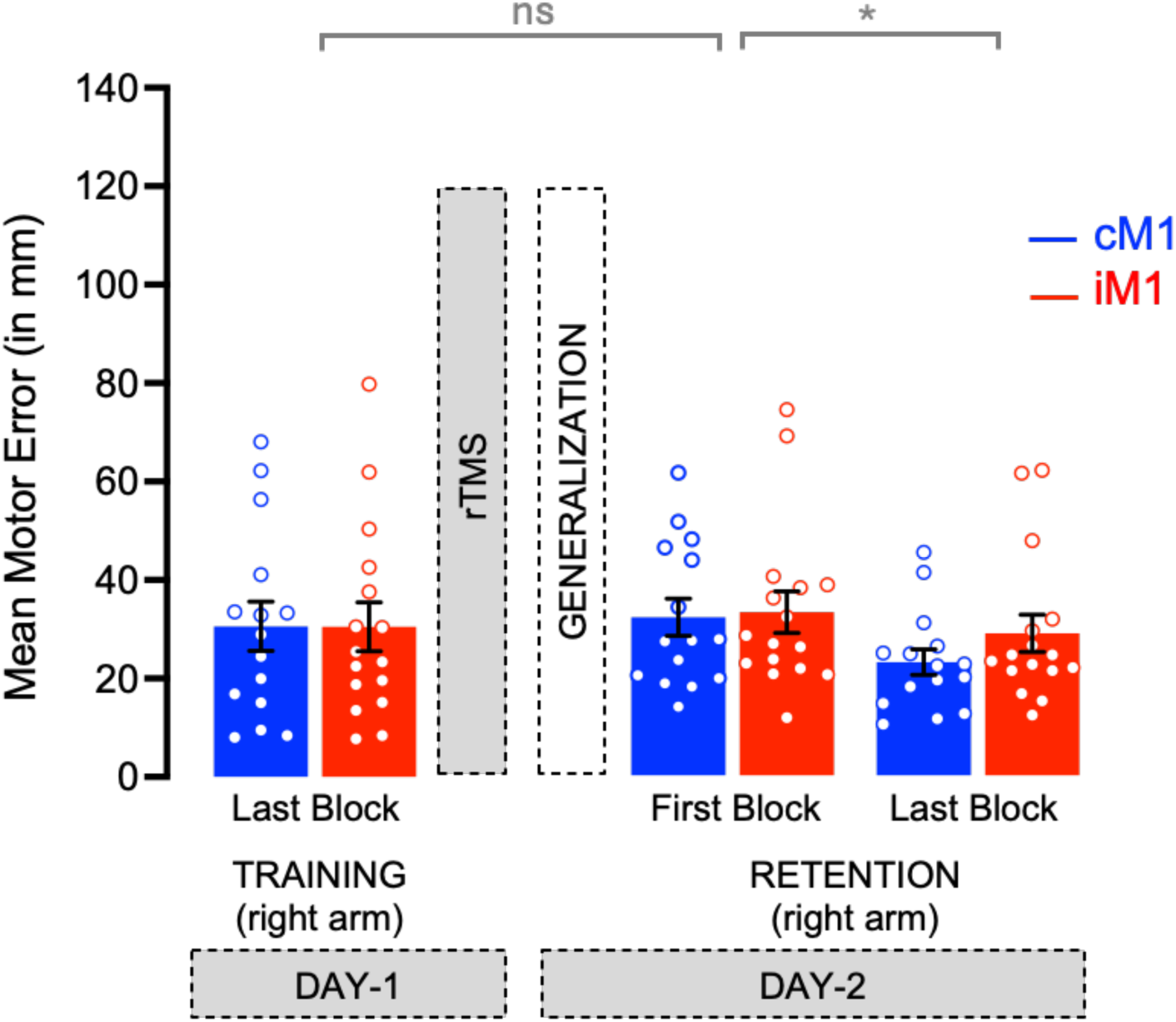
Intramanual Skill Retention. Left Panel: Mean motor error in last training block on Day-1. Right Panel: Same for first and last retention blocks on Day-2. Note the similar amount of motor errors when comparing last training and first retention blocks, in both the cM1 (in blue, n=15) and iM1 (in red, n=16) groups, suggesting similar level of skill retention across two days in both groups. Moreover, both groups further improved performance of their right trained arm over the two blocks of this session on Day-2 with no group differences. Error bars on the figure denote SE. Colored circles denote individual participants.

Taken together, our behavioural results show that applying 1Hz rTMS over contralateral M1, as compared to ipsilateral M1 at the end of skill training, led to reduced interlimb generalization of the newly acquired skill to the untrained arm, without affecting the subsequent retention performance of the trained arm.

### 3.2 Motor Neurophysiology

#### Learning induced changes in CSE

We were curious to know whether learning on our skill task induced excitability changes in the primary motor cortex, as reported in previous research using different motor tasks (Uehara et al., 2018; Spampinato et al., 2019; Yadav et al., 2025b). We explored whether CSE changed within contralateral and ipsilateral M1 due to skill training in our study by comparing the MEPs obtained in right FDI of cM1 and left FDI of iM1 groups, before and after the training session (see Figure 5A). Our analysis revealed no significant group (F_1,29_ = 0.3557, p=0.5555, η^2^_p_=0.0121), timing (F_1,29_ = 0.3093, p=0.5824, η^2^_p_=0.0105) or group X timing interaction (F_1,29_ = 2.6737, p=0.1128, η^2^_p_=0.0844).

**Figure 5.**
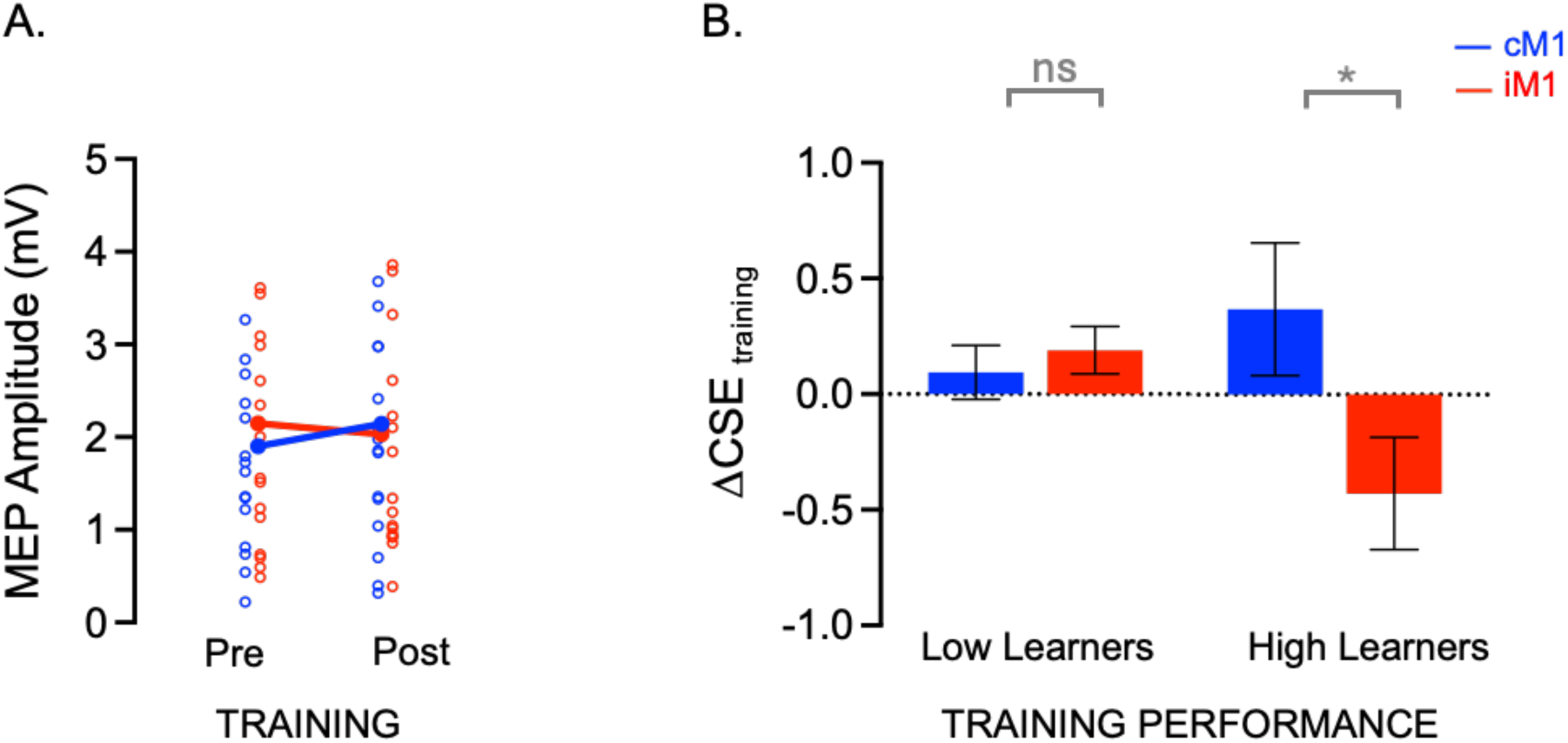
CSE changes related to skill training. **A) Pre- and Post-training MEP amplitudes.** No significant change in CSE as a result of training in the two groups. **B) CSE Changes in Low and High Learners of cM1 and iM1 groups.** A median-split based on learning revealed significant differences in training-related changes in CSE (τ<CSE_training_: post-training minus pre-training) between cM1 and iM1, only in high learners of the two groups, but not for low learners. Error bars on the figure denote SE. Colored circles denote individual participants.

However, upon performing some exploratory analyses, we noticed heterogeneity in the amount of learning across subjects. We, therefore, categorized subjects into “High” and “Low” learners based on group wise median-split of their training motor error data (first training – last training blocks) to evaluate any effect of learner-type on CSE changes induced by training (ΔCSE_training_: Post-Training – Pre-Training MEPs). Our two-way ANOVA on this ΔCSE_training_ with group (cM1, iM1) and learner-type (High, Low) as factors revealed a significant group X learner-type interaction (F_1,27_ = 4.5834, p=0.0415, η^2^_p_=0.1451). The Tukey post hoc test showed a significant difference in ΔCSE for “High” learners of the two groups (p=0.0485; ΔCSE for cM1: 0.36 ± 0.20 mV and iM1: -0.42 ± 0.20 mV). These results suggest that in high-learners, skill training resulted in a larger increase in CSE (i.e., higher Post-Training MEPs as compared to Pre-Training MEPs) when measured on the trained side (TMS over contralateral M1) than on the untrained side (TMS over ipsilateral M1; Figure 5B). On the other hand, such difference was not found for low-learners (p=0.9885 [ΔCSE for cM1: 0.09 ± 0.21 mV and iM1: 0.18 ± 0.20 mV). These results suggest that learning induces greater increases in CSE on the trained side than on the untrained one, but this effect is only apparent in individuals showing greater learning. However, please note that these analyses were conducted on small samples and should therefore be considered preliminary, aiming to generate hypotheses and guide future investigations rather than establish strong conclusions.

#### rTMS related changes in CSE

Finally, related to the 1 Hz rTMS intervention used in our study, we examined MEPs using a two-way ANOVA with group (cM1, iM1) and rTMS timing (Pre, 0 min, 5 min, 10 min) as factors. This analysis revealed a significant effect of rTMS timing (F_3,29_ = 3.3029, p=0.0240, η^2^_p_=0.1022), but no group (F_1,29_ = 0.0078, p=0.9299, η^2^_p_=0.0000) or timing X group interaction (F_3,29_ = 0.4285, p=0.7331, η^2^_p_=0.0145). Surprisingly, related to the rTMS timing effect, the post-hoc test revealed a significant increase in MEP amplitudes between Pre and the Post-rTMS 10 min timing (p=0.0490 [Pre-rTMS: 2.08 ± 0.11 mV and Post-rTMS 10min: 2.50 ± 0.11 mV]). This suggests that, irrespective of whether the 1Hz rTMS was applied on (left) cM1 or (right) iM1, it produced a significant increase in MEP amplitudes at 10 min following the end of the neuromodulation intervention. Hence here, rTMS seems to have increased rather than decreased CSE associated with the stimulated M1 (see Figure 6A). This finding of ours contrasts with the anticipated inhibition effect based on past research (Wassermann et al., 1996; Chen et al., 1997, Touge et al., 2001; Fitzgerald et al., 2002; Chouinard et al., 2003; Lang et al., 2006). Instead, our data adds to the literature that is suggestive of mixed findings including no change in CSE or facilitation especially at higher intensities which we discuss later in the discussion section (Pascual-Leone et al., 1994; Siebner et al., 1999; Siebner et al., 2004; Fitzgerald et al., 2006; Lang et al., 2006; Hoogendam et al., 2010; Veldema et al., 2023).

**Figure 6.**
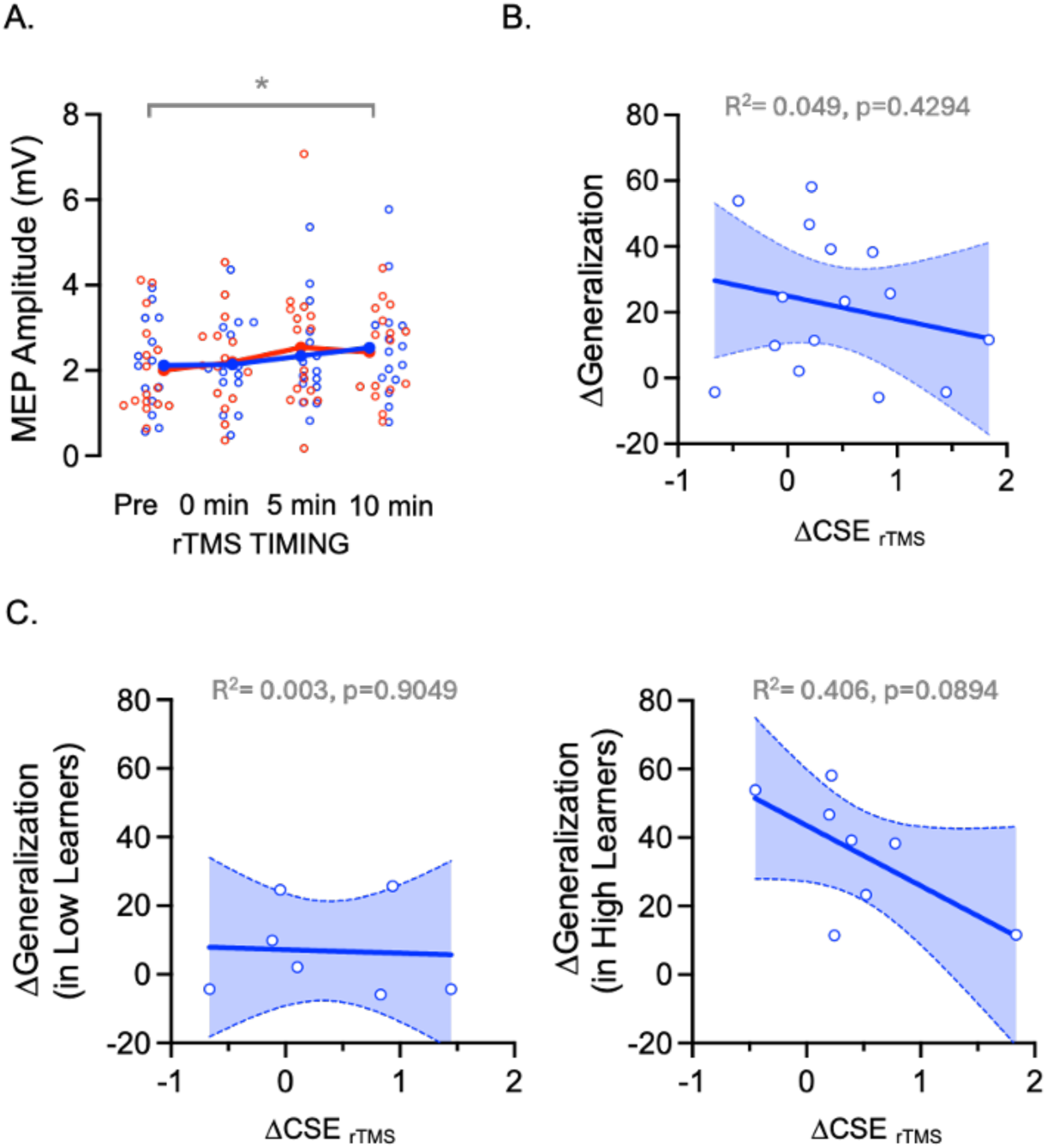
CSE changes related to the rTMS intervention. **A) MEP amplitudes across rTMS timings.** rTMS induced a significant increase in MEP amplitudes (mV) for Post-rTMS 10 min compared to Pre-rTMS, irrespective of group type. **B. Correlation with Generalization in cM1 group.** For cM1 group (blue in A), who exhibited reduced interlimb skill generalization between first training and first generalization blocks (τ<Generalization: lower values mean lower generalization), no significant correlation was found between their generalization performance and rTMS induced CSE changes (τ<CSE _rTMS_: Post-rTMS 10 min compared to Pre). **C. Same correlation in Low- (left panel) and High Learners (right panel) of cM1 group.** A near significant correlation was noted for the high learners of this group, but not for the low learners. These results are of course to be taken with caution given the small size of each subgroup. Error bands on the figure denote SE. Colored circles denote individual participants.

With significant CSE facilitation observed at 10 min following rTMS, we wanted to examine whether there was any correlation between the strength of this CSE increase and the reduced amount of interlimb generalization observed in cM1 group in our study. To do this, we performed a linear correlation between the rTMS-related CSE change (ΔCSE_rTMS_ = Post-rTMS 10min – Pre-rTMS) and interlimb generalization performance (ΔGeneralization = Motor Errors on First Training block – First Generalization block) for the cM1 group (Figure 6B). Our results revealed no significant correlation between ΔCSE_rTMS_ and ΔGeneralization (R^2^=0.049, p=0.4294). Further, upon performing an exploratory analysis on this correlation based on learner-type (high-versus low-learners as described earlier), we noted a marginally significant correlation between ΔCSE_rTMS_ and ΔGeneralization for high-learners of the cM1 group (R^2^=0.406, p=0.0894) with no significant correlation for low-learners (R^2^=0.003, p=0.9049) (see Figure 6C). These preliminary findings suggest that the rTMS-induced facilitation of cM1 may have negatively affected generalization in high learners: the stronger the facilitation, the weaker the generalization. Our observation here supports the notion the rTMS can induce opposite effects on motor behavior and excitability changes (Hoogendam et al., 2010; Veldema et al., 2023).

## 4. DISCUSSION

In this study we primarily set out to probe the causal role of contralateral primary motor cortex (cM1) in intermanual skill generalization by testing the untrained non-dominant left arm at 24 hours following right hand training. As hypothesized, we found that non-invasively stimulating cM1 (versus iM1) using rTMS intervention did impair skill generalization to the untrained arm. Despite this reduction in the amount of generalization in the cM1 group, compared to iM1 group, overall, both groups still benefitted from prior training: the naïve non-dominant untrained (left) arm performance on Day-2 was significantly better than that of the naïve trained (right) arm on Day-1. And contrary to our hypothesis, rTMS to cM1 in our study did not impact intralimb skill retention of the trained arm which was tested at the end of Day-2. Behaviorally, these findings suggest that intermanual generalization of skill memory following de novo motor learning can be hampered if cM1 is targeted immediately following skill learning. Next, we noted that in the subset of participants who exhibited higher learning, CSE changes measured in the trained hand (assessed with cM1 TMS) were larger (i.e., an increase in training-induced MEPs) as compared to CSE changes in untrained hand (assessed with iM1 TMS). Finally, and somewhat unexpectedly, our rTMS intervention produced a facilitation of CSE, which tended to correlate negatively with the degree of intermanual generalization on Day-2 in high-learners of cM1 group: the more cM1 rTMS induced CSE facilitation on Day-1, the lower the intermanual generalization on Day-2.

Our current findings build upon our prior work in which we investigated whether it is possible for the untrained arm to access newly acquired fine motor skill memory from training of the other arm; and if yes, then what are the underlying mechanisms driving this phenomenon in young healthy individuals. We have previously reported (Yadav and Mutha, 2020; Yadav et al., 2025a) that the phenomenon of intermanual skill generalization to the untrained arm- (1) is robust following successful acquisition of novel skill memory representations, and (2) is negatively impacted when individuals engaged in a secondary task immediately following training, suggesting that post-training skill memory consolidation/stabilization processes operating endogenously also impact the amount of generalization. In this study, we took the next obvious step to explore the neural substrate involved in mediating these processes and focused on cM1- a key brain region for skill learning and consolidation (Floyer-Lea and Matthews, 2005; Doyon et al., 2009; Narayana et al., 2014; Kawai et al., 2015; Gerardin et al., 2024), specifically in the context of predictable skill reaching movements (Kantak et al., 2010a). Thus, we non-invasively stimulated cM1 (with iM1 serving as an active control group) immediately after learning with the aim to perturb the newly acquired skill memory and found that it negatively affected the untrained left arm performance on the next day. However, the overall skill performance of the untrained arm in both groups was significantly better as compared to the naïve training arm (when comparing first generalization and first training blocks) thereby suggesting that the untrained arm still benefitted from the trained arm learning despite the stimulation intervention. Moreover, the untrained arms further learned on Day-2 and reached the pre-stimulation performance level of the trained arm at the end of Day-1. Additionally, perturbing the newly acquired memory with cM1 (or iM1) stimulation did not impact intralimb retention: trained arm tested on Day-2 maintained its pre-stimulation performance levels as well. This intact retention could possibly be attributed to our study design which allowed the right trained arm to not only benefit from its original training on Day-1, but also from the left untrained arm test performance that occurred just before on Day-2, thus resulting in another instance of intermanual skill generalization (left to right arm in this case). Thus, although stimulation of cM1 reduced skill generalization to the untrained arm, it did not prevent further learning in this arm or impair subsequent test performance of the trained arm on Day-2. In other words, although cM1 neuromodulation weakened generalization, it did not prevent some generalization from occurring. These observations could imply the following: (1) involvement of additional brain regions that participate in skill learning and memory consolidation and thereby impact future motor skill performances such as generalization, (2) 1Hz rTMS intervention could not efficiently perturb the offline memory consolidation processes operating within cM1 (as shown by Psurek et al., 2021 for motor sequence learning), and (3) variability observed in the amount of skill learning on our task and differential rTMS effects on high versus low learners. Based on our behavioral and neurophysiology data, we have reflected on these points in the coming section along with the limitations posed by a non-invasive brain stimulation approach such as rTMS for probing human skill behavior.

First, we would like to understand the underlying neural bases for the skill behavior observed in our study. We found that three main frameworks can be identified from the broader motor learning literature that may help us uncover how sharing or transfer of movement related information in the brain and between limbs (such as arms) facilitates motor behavior. The first framework for cross-limb transfer of learned movements is referred to as ‘*callosal access hypothesis’*, and according to this, skill memory representations formed following learning reside in the trained contralateral hemisphere which in turn can be accessed by the untrained hemisphere/arm system through corpus callosum (Taylor and Heilman, 1980; Anguera et al., 2007, Ruddy and Carson, 2013). Second framework for intermanual transfer of motor learning is known as ‘*cross-activation hypothesis*’ (Parlow and Kinsbourne, 1989; Ruddy and Carson, 2013) which posits that extensive unilateral training generates cortical activity not only in the contralateral hemisphere but also the ipsilateral hemisphere. This ipsilateral activity in turn facilitates performance benefits to the untrained arm (Anguera et al., 2007; Lee and Carroll, 2007). Lee et al., (2010), for instance, showed that rTMS to the untrained hemisphere reduced the amount of intermanual generalization of ballistic movements, providing support to the view that cortical processes in the hemisphere ipsilateral to the trained arm can contribute to transfer.

In our case, however, rTMS to cM1, and not iM1 led to an impairment in skill generalization to the untrained arm, thus favoring the *callosal access hypothesis* as opposed to the *cross-activation hypothesis*. However, our findings also show that the naïve untrained arm tested on Day-2 still benefitted from the training arm even after rTMS intervention. This indicates that skill learning and memory consolidation related neuroplastic changes that could drive such generalization are possibly not strictly restricted to cM1. Consistently, a number of neuroimaging findings have highlighted the involvement of a wide number of brain regions in addition to M1, such as the somatosensory cortices, SMA, DLPFC, posterior parietal cortex, basal ganglia and cerebellum that participate in skill learning (Hikosaka et al., 2002; Doyon et al., 2002; Floyer-Lea and Matthews, 2005; Doyon and Benali, 2005; Dayan and Cohen, 2011; Vassiliadis et al., 2024). The seat(s) of skill memory representation in the brain, thus, has a lot of implications for skill generalization. This brings us to a third framework, namely the ‘*bilateral access hypothesis’* (which Ruddy and Carson, 2013 have merged together with the ‘callosal access’ framework). This model of intermanual generalization acknowledges that learning related neuroplastic changes not only involve cM1, as implied by the callosal access hypothesis, but also additional brain areas that participate in skill learning and memory consolidation. Skill representation developing or consolidating in these additional brain regions, in turn, can be bilaterally accessed by the untrained or trained hemisphere/arm whenever the task is performed again. Our current data with overall significant skill transfer (though reduced in the group that received cM1 stimulation) as well as intact intralimb retention lend support to this framework hinting participation of additional brain regions. Future work targeting some of these above-mentioned areas involved in skill learning (such as DLPFC or SMA that could potentially be involved in skill task such as ours) can help elucidate regions in addition to cM1 that are facilitating skill generalization and retention.

Finally, we would like to briefly comment about the role of rTMS in our study and its variable effects. First, we noted a facilitatory effect of rTMS on CSE in both our groups as opposed to the traditionally expected inhibitory effect induced by low-frequency stimulation (1 Hz) reported earlier (Chen et al., 1997; Boroojerdi et al., 2001; Fitzgerald et al., 2006; Kantak et al., 2010a, 2010b; Psurek et al., 2021). Our findings, on the other hand, are consistent with other work who report mixed effects of rTMS on excitability (Pascual-Leone et al., 1994; Siebner et al., 1999; Fitzgerald et al., 2006; Lang et al., 2006, Hoogendam et al., 2010; Siebner et al., 2022; Veldema et al., 2023). Taken together this shows heterogeneity in how rTMS modulates underlying cortical activity. Furthermore, our data show an interesting effect with cM1 rTMS facilitating CSE while impairing generalization, an effect that may be specific to high learners, based on our preliminary results. This shows a dissociation between TMS induced underlying excitability changes and its effect on behavior, which has been reported by other as well (Jäncke et al., 2004; Schramm et al., 2019; Kim et al., 2021; Psurek et al., 2021; Veldema et al., 2023). We think such variability seen in studies investigating neurophysiology measures and behavior can be attributed to many factors. These include differences in stimulation parameters across studies (Maeda et al., 2000; Kantak et al., 2010b), such as the intensity of stimulation (we used 110% of rMT for rTMS as compared to a subthreshold intensity of 90%) and its duration (we applied 1800 pulses, a maximum deemed safe for efficient brain stimulation). The type of machine and coil used could also matter (Lang et al., 2006). Moreover, timing of measurement (such as at 0, 5, 10 mins post-stimulation) and endogenous brain state while applying rTMS (i.e., different learning levels) could also influence the pattern of findings. Each of these factors requires a separate detailed investigation with larger sample size in order to fully establish the efficacy of this non-invasive brain stimulation procedure for studying human behavior. In summary, our current study employs rTMS to elucidate the role of cM1 in human skill generalization of de novo learning and extends previous literature showing its involvement in human motor learning, plasticity and memory consolidation.

## Acknowledgments

We would like to thank Bhoomika Sonane for her assistance with behavioral task setup, and Julian Lambert and Benvenuto Jacob for their technical assistance with setting up the recording devices used in this study.

